# Measles Whole Genome Sequencing by an Illumina Tiled Amplification Method

**DOI:** 10.64898/2026.05.13.724913

**Authors:** V Zubach, S Ashfaq, S Van Driel, B Kaplen, G Peters, V Laminman, A Go, C Bonner, M Graham, J Hiebert

## Abstract

Measles virus remains a significant global health threat, and despite the availability of an effective vaccine, measles cases continue to increase worldwide in recent years. Genomic surveillance has become an essential tool for monitoring virus circulation and investigating outbreaks. Here, we describe a wet-laboratory method for whole-genome sequencing of measles virus using a tiled amplicon approach and Illumina sequencing technology. A previously published Oxford Nanopore–based tiled primer scheme was adapted to include both circulating measles genotypes and for use on the Illumina platform. Two Illumina library preparation kits, Illumina DNA Prep (IDP) and Nextera XT (XT), were evaluated for performance. The IDP kit demonstrated more complete genomes and consistent genome coverage compared with XT. Using quantified reference genomes, the limit of detection was determined to be 10,000 genome copies for genotype B3 and D8. Sequence accuracy was evaluated using previously characterized clinical samples and showed high concordance. This method provides a reliable and sensitive approach for measles virus whole-genome sequencing using Illumina platforms and is suitable for genomic surveillance applications.

## Introduction

Measles virus (MeV) is a highly contagious respiratory pathogen and remains a major threat to global public health. Although widespread use of an effective live-attenuated vaccine has substantially reduced global measles incidence, recent years have seen a marked resurgence of measles cases across multiple regions, largely driven by suboptimal vaccination coverage, immunization program disruptions, and increased population mobility (1). These trends threaten measles elimination goals and highlight the need for enhanced surveillance strategies (2).

Molecular surveillance plays a central role in measles monitoring by enabling the identification of circulating MeV genotypes and tracking chains of transmission. The World Health Organization currently recommends MeV genotyping based on sequencing a 450-nucleotide region of the nucleoprotein gene (N450), which has historically been sufficient to distinguish between globally circulating measles virus genotypes (3). However, the progressive reduction in measles genetic diversity, with only B3 and D8 genotypes now detected worldwide, has limited the discriminatory power of partial-genome sequencing for outbreak investigation and transmission analysis (4).

Whole-genome sequencing (WGS) provides higher resolution than targeted sequencing and has proven valuable for resolving complex transmission networks, distinguishing between endemic circulation and repeated importation events, and supporting elimination efforts in low-incidence settings (5). Despite these advantages, routine implementation of MeV WGS has been limited by technical challenges, including low viral loads in clinical specimens and the need for cost-effective, sensitive laboratory workflows suitable for public health laboratories (6).

One of the main technical challenges in MeV genome sequencing is obtaining good sequencing coverage throughout the long non-coding region between the matrix and fusion genes (MF-NCR). This region proves difficult to sequence due to its high GC content, frequent homopolymer stretches, and lower abundance compared to coding regions. The MF-NCR has attracted considerable interest to sequence because of its known sequence variability, which can aid in molecular resolution and surveillance. The MF-NCR is also a known hotspot for insertions and deletions, leading to variation in genome lengths (7). Tiled amplicon sequencing approaches have emerged as an effective solution for viral WGS, offering high sensitivity, uniform genome coverage, and compatibility with samples containing limited amounts of viral RNA. This strategy has been widely adopted for a range of RNA viruses and is particularly well-suited for outbreak-driven genomic surveillance where rapid turnaround and scalability are essential (8,9). A tiled primer scheme for measles virus WGS has previously been described using Oxford Nanopore Technologies (ONT) (10); however, differences in sequencing chemistry, read length, and library preparation requirements necessitate adaptation and validation for Illumina sequencing platforms, which remain the most commonly used platforms in public health genomics laboratories (11). Another popular tiled primer scheme for measles is the ARTIC network’s 400bp scheme, which amplifies the complete MeV genome in 2 primer pools (12).Based on experience with the development of a tiled amplicon scheme for ONT (10), we have found that the use of six primer pools enables uniform amplification and allows enhanced amplification performance across difficult-to-sequence regions of the measles virus genome.

In this study, we developed and evaluated an Illumina-based tiled amplicon approach for MeV WGS derived from the previously published method (10), and specific assessment of two Illumina library preparation kits. We determine the analytical sensitivity and accuracy of the final method using reference materials and previously characterized clinical specimens.

## Methods

### Reference Genomes and Samples

Genotype D8 (VR-1980) and B3 (VR-1981) MeV isolates with downloadable WGS data were purchased from American Type Culture Collection (ATCC). Extracted nucleic acid was quantified with a QuantStudio Absolute Q Digital PCR (dPCR) system (Applied Biosystems) using primers targeting the measles nucleoprotein gene and a 10-fold dilution series from 10^5^ to 100 copies/µL was prepared as previously described (10).

Archival MeV RT-PCR-positive clinical specimens (n = 64) that were referred to the Public Health Agency of Canada’s (PHAC) National Microbiology Laboratory (NML) for the purpose of routine measles genotype surveillance were included. Research ethics board review was not required as per Article 2.5 of the Tri-Council Policy Statement: Ethical Conduct for Research Involving Humans – TCPS 2 (2022) (13). The genotype distribution was 46 B3 and 18 D8. For 35 of the samples (27 genotype B3 and 8 genotype D8), WGS were previously obtained by a hybrid capture method (14) and used to determine sequence accuracy.

### Primer design

Primer design was based on the tiled amplicon scheme described by Zubach et al. 2025 (10), which included the addition of supplemental primers specifically targeting the MF-NCR to ensure complete and consistent coverage breadth across the measles virus genome. To improve amplification performance in the genomically challenging MF-NCR and reduce primer–primer interactions, primers were distributed across six distinct primer pools. The original primer set developed by Zubach et al. 2025 (10) was specific to measles virus genotype D8; selected primers were therefore modified to support amplification of both genotype D8 and B3 strains. Amplicon sizes range from 341 to 1011 base pairs. Primer sequences, pool assignments, concentrations, and expected amplicon sizes are provided in Supplementary Table S1.

### Extraction, PCR and Library Preparation

Total nucleic acids were extracted using the MagNA Pure 96 DNA and Viral NA large volume kit (200 µL of specimen) on the MagNA pure 96 (Roche Diagnostics catalog 06374891001) or with the QIAamp Viral RNA Mini Kit (280 µL of measles viral isolate) (QIAGEN catalog 52904) and eluted in 50 µL, following manufacturer’s instructions. Extracted RNA was stored at −80°C if not used immediately. Reverse-transcription was performed with 7.5 µL of extracted total nucleic acid using Superscript IV (ThermoFisher Scientific catalog 18090200) according to the manufacturer’s instructions but included betaine (Sigma-Aldrich catalog B0300) to a final concentration of 0.2 M in a 30 µL reaction. Multiplex PCR occurred when 2 µL of the prepared cDNA was amplified in six different pools using Q5 HotStart High Fidelity 2× mastermix (New England BioLabs catalog M0494). For primer sequences, concentrations and pools see Supplementary Table S1. Multiplex PCR reactions were carried out in 12.5 µL according to the manufacturer’s guidelines. Initial denaturation took place at 98°C for 30 s, amplification followed with 34 cycles of denaturation at 98°C for 15 s, and primer annealing/extension at 68°C for 5 min. After PCR amplification, amplicons from the six pools were combined and purified using PCR Clean DX beads (Aline Biosciences, catalog C-1003-450), at a 1.8x bead ratio, then eluted in 0.01M Tris pH 8.0. The DNA was then quantified on the Qubit Flex (Life Technologies) or FilterMax F5 (Molecular Devices) using the 1X dsDNA Broad Range Assay Kit (Life Technologies catalog Q33266).

Illumina Nextera XT (Illumina catalog FC-131-1096) and Illumina DNA Prep (Illumina catalog 20060059) libraries were prepared following an in-house optimized reduced-volume protocol. Libraries were indexed with 10 bp unique dual indexes (UDIs) from Integrated DNA Technologies. Final libraries were quantitated using the PicoGreen assay (Life Technologies catalog P7581) on the FilterMax F5 and then equimolarly pooled. Pools were size selected for 150-1000 bp on the Blue Pippin Instrument (Sage Science) using 1.5% Agarose Pippin Cassette (D-Mark Biosciences catalog SAG-BDF1510). Final pools were assessed on the Agilent TapeStation 4200 using the Genomic DNA ScreenTape (Agilent Technologies catalog 5067-5365) to determine average fragment size and quantitated via Qubit 1X dsDNA High Sensitivity Assay Kit (Life Technologies catalog Q33231). Paired-end (2×300 bp) sequencing was performed on the Illumina NextSeq 1000/2000 platforms using XLEAP-SBS chemistry. Upon sequencing run completion, fastq files were generated using BCL Convert. See Figure 1. for an overview of the complete Illumina Tiled Amplification workflow.

**Figure 1.**
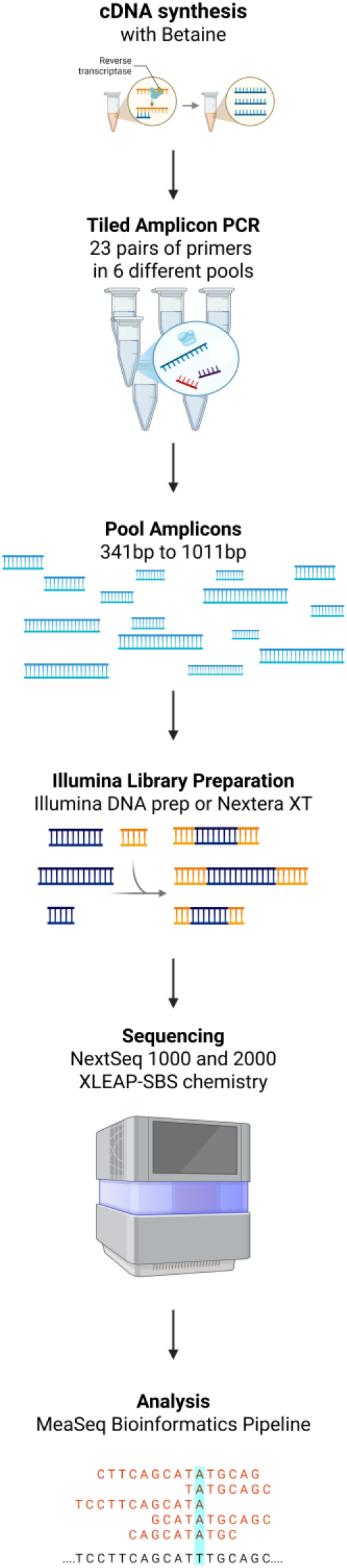
Illumina tiled amplicon workflow. Schematic overview of the amplicon tiling strategy used for WGS of measles virus on the Illumina platform. Ribonucleic acid (RNA) from clinical specimens is reverse-transcribed in the presence of betaine to generate cDNA. Six primer pools, comprising 23 primer pairs, are prepared and used in multiplex PCR reactions to generate tiled amplicons spanning the viral genome. Amplicons range in size from 341 to 1,011 bp. Amplicons from all pools are combined and purified, followed by library preparation using the IDP or XT kits. Prepared libraries are loaded onto the Illumina NextSeq 1000/2000 sequencer, and sequencing data are subsequently generated for downstream analysis using the MeaSeq pipeline.

### Bioinformatic Analysis

Post sequencing quality control and consensus sequence generation were completed using the measles specific bioinformatics pipeline MeaSeq (https://github.com/phac-nml/measeq) (15). For consistency across analyses, all WGS consensus sequences were trimmed to exclude genomic termini (WGS-t), beginning at the start codon of the nucleoprotein open reading frame (ORF) and ending at the stop codon of the large ORF. For accuracy assessment, samples had to have consensus sequences achieved with a minimum of 10× coverage across the genome. Trimmed whole genome consensus sequences were 15,678 or 15,684 nucleotides in length. Sequences flagged by MeaSeq for indels underwent minor manual curation.

## Results

### More complete genomes achieved with the Illumina IDP kit than the XT) kit

A total of 83 sequences were generated from 64 unique samples, including serial dilutions from 4 different specimens. The median Crossing Point (CP) value was 26.74 and included dilutions beyond a value of 30 which is the typical threshold for attempting whole genome sequencing (range of 18.63 to 33.95) (16). The sequences had varying levels of genome completeness and were compared across the IDP and XT library preparation kits. Metrics assessed were number of sequences yielding complete genomes, average genome completeness, and median read depth by each library preparation kit. Median sequencing depth for the IDP and XT kits was 633.52 and 771.98 respectively. Dropout regions were frequently observed in libraries prepared with the XT kit (Figure 2), most commonly affecting MVS7 and MVS8, which span the MF-NCR and are 505 bp and 765 bp in length, respectively. Average sequence coverage completeness across the 83 genomes, expressed as a percentage of the genome, was only slightly higher with the IDP kit (96.85%) compared to the XT kit (96.36%). However, additional dropouts observed with the XT kit resulted in fewer number of complete genomes, defined as 100% sequence coverage across the WGS-t, with only 36.1% of samples meeting this threshold, compared to 55.4% for the IDP kit. Supplementary figure 2 provides a coverage depth plot comparing a sample prepared with the IDP kit versus the XT kit. Overall, the Illumina DNA Prep kit produced more complete genomes with more consistent coverage and was therefore selected for the final workflow.

**Figure 2.**
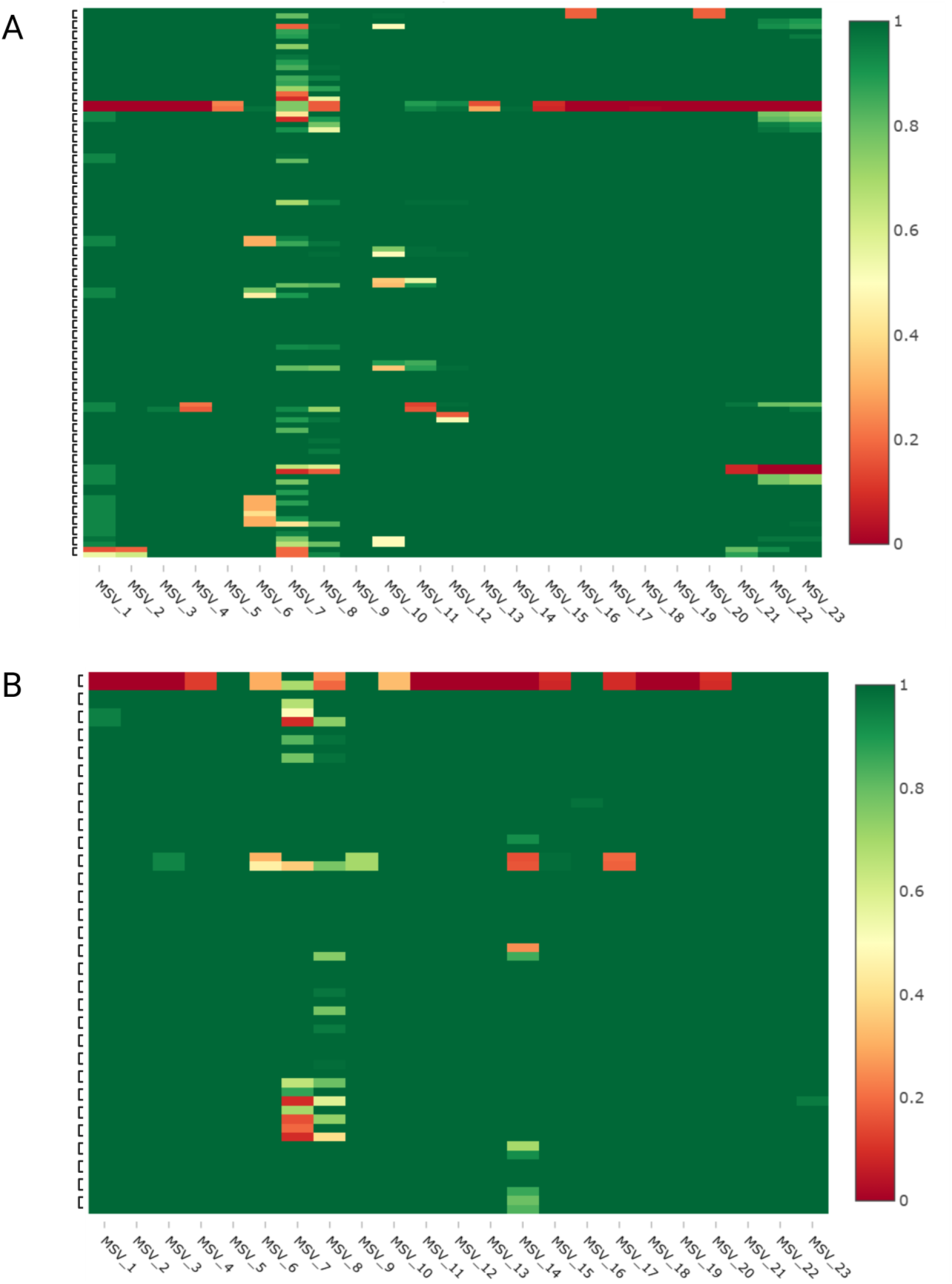
Amplicon completeness heatmap across the genome comparing IDP and Nextera XT library preparations. (A) Genotype B3 samples (B) Genotype D8 samples (n=83). Samples are shown in matched pairs, with the IDP prepared sample displayed above the corresponding XT prepared sample. Refer to Supplementary Table 1 for the primers associated with each amplicon.

### Limit of Detection and Accuracy using the Illumina DNA Prep Kit and the ATCC reference genomes

Serial dilutions of quantified B3 and D8 ATCC reference MeV genomes were sequenced three times in three separate library preps and sequencer runs with the IDP kit. The CP values for these serial dilutions ranged from 25.00 to 32.87 for B3 and 25.00 to 33.95 for D8, exceeding the typical CP value cut-off of 30 used for selecting samples for WGS. Although both genotypes produced sequences that were complete down to 100 copies/µl, genotypes B3 and D8 consistently yielded complete genomes at 10,000 copies/µl (Table 1). Median depth for B3 and D8 began to drop at 100 copies/µl. All serial dilutions for B3 genotype yielded 100% accurate sequences compared to the ATCC supplied reference genome. For genotype D8, all serial dilutions but the 100 copies/ul had 100% accurate sequences. One of the replicates for the 100 copies/ul dilution had a 100% accurate sequence, while 1 sequence had 2 ambiguous mismatches and the other sequence had 1 ambiguous mismatch and 1 nucleotide change.

**Table 1.**
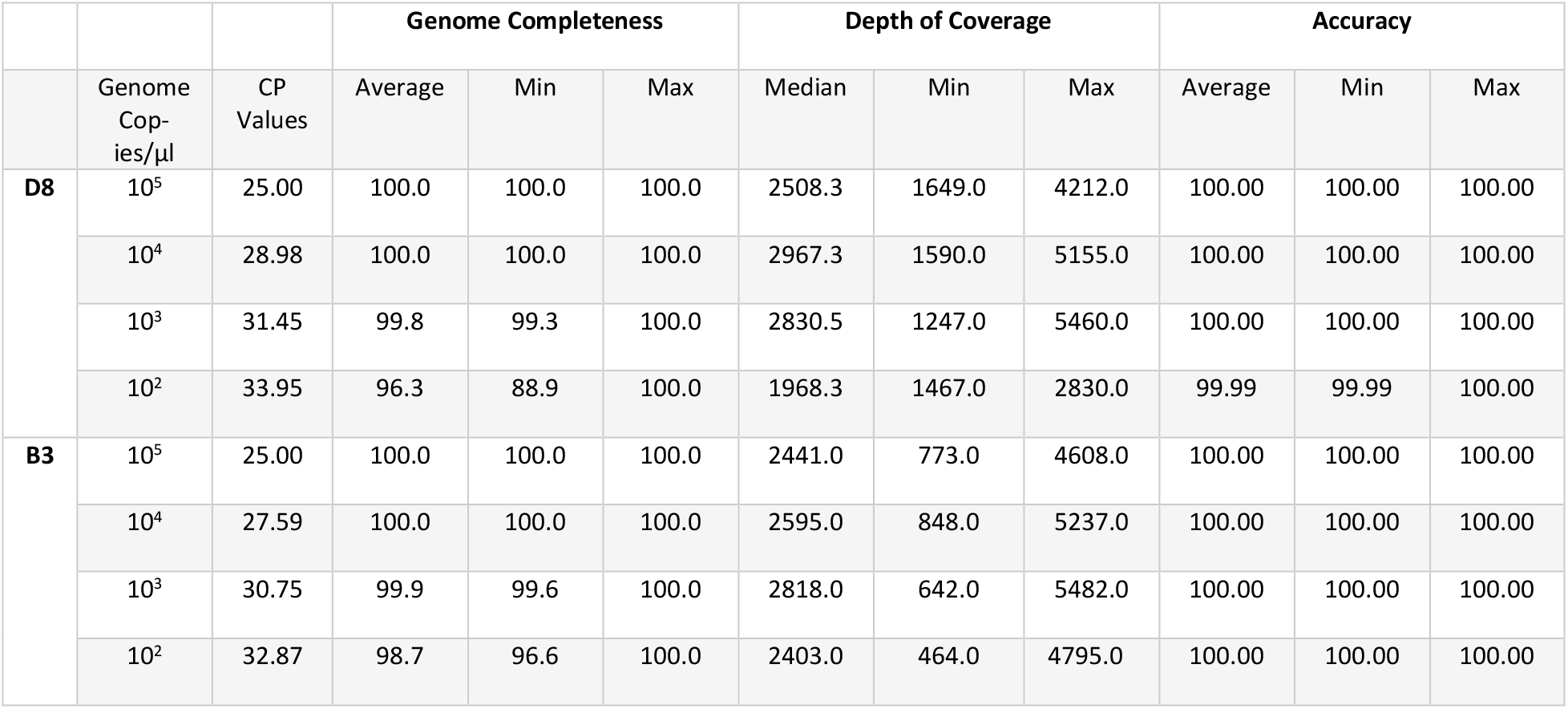
Limit of detection, Median depth and Accuracy across the serial dilution for the ATCC’s MeV reference genomes. Each dilution was sequenced three times.

### Complete concordance achieved when IDP WGS compared to previously characterized WGS

Thirty-five clinical samples, including 27 genotype B3 and 8 genotype D8 specimens sequenced using the pooled tiled amplicon approach, were compared with available WGS generated using an Illumina hybrid capture method (14). Crossing point (Cp) values for these samples ranged from 18.63 to 31.49. Consensus genome lengths for the sequenced samples were either 15,678 or 15,684 base pairs.

Thirty-two of the 35 samples showed complete (100%) concordance between the two sequencing approaches. Three samples each contained a single heterogeneous site at distinct genomic positions; two of these heterogeneities were detected by both the tiled amplicon and hybrid capture methods. In the remaining sample, a heterogeneous site identified in the hybrid-capture data was resolved as a single matching nucleotide in the tiled amplicon consensus sequence. One additional sample contained three ambiguous nucleotides at closely spaced but distinct genomic positions. Inspection of the corresponding BAM files indicated the presence of low-quality reads contributing to these ambiguities. Removal of the problematic reads would result in a corrected WGS consensus sequence that was fully concordant with the reference sequence.

## Discussion

In this study, we describe and evaluate an Illumina-based tiled amplicon workflow for WGS of MeV that was adapted from a previously published Nanopore-based approach (10) and expanded to include both currently circulating genotypes, B3 and D8. Our results demonstrate that this method reliably produces accurate whole-genome sequences from clinical specimens and reference materials and represents a practical option for measles virus genomic surveillance using Illumina platforms.

Comparison of library preparation kits revealed that the IDP outperformed XT kit, yielding a higher proportion of complete genomes and more uniform genome coverage. Although median sequencing depth was comparable between the two kits, XT libraries exhibited a greater number of dropout regions especially in the MF-NCR, consistent with the known sequence and GC biases associated with transposase-based fragmentation and low-input PCR normalization, whereas IDP employs bead-linked transposomes and reduced amplification bias (17). These findings support the use of IDP for tiled amplicon measles virus sequencing.

Analytical sensitivity for measles virus genotypes B3 and D8 consistently yielded complete genome sequences at template concentrations of 10,000 copies/µL, corresponding to CP values of 27.59 and 28.98, respectively. For both reference genomes, individual replicates were able to achieve complete genome coverage at concentrations as low as 100 copies/µL. Nevertheless, complete genomes for both genotypes were reproducibly obtained at virus concentrations consistent with those typically observed in routine diagnostic specimens, supporting the suitability of this approach for genomic surveillance applications.

Using 10-fold serial dilutions of genotype B3 and D8 reference genomes from 100 to 100,000 copies/µl, the consensus sequence accuracy was 100% across all concentrations of genotype B3 genomes. For genotype D8, two of the three consensus sequences generated using a 100 copies/µL dilution 100% accurate. However, the CP value at this concentration was 33.95, exceeding the threshold of 30 that is commonly used for selecting samples for WGS.

Sequencing of 35 clinical samples produced 31 consensus sequences that were fully concordant with the previously established whole-genome sequences generated using an Illumina hybrid-capture method (14). The detection of heterogeneous sites by both sequencing approaches in two samples further supports the reliability of the tiled amplicon workflow and suggests that observed base-call ambiguities likely reflect genuine biological variation rather than methodological artefacts. In one instance, a heterogeneous site identified by the hybrid-capture approach generated a non-ambiguous nucleotide assignment in the tiled amplicon consensus sequence. In another sample, three closely spaced but distinct ambiguous nucleotide positions were observed relative to the reference sequence; however, inspection of the corresponding BAM files indicated that low-quality reads were contributing to these calls. Removal of the affected reads would result in a consensus sequence fully concordant with the reference. The presence of two distinct whole-genome sequence lengths within the sample set, attributable to a six-base-pair insertion in the MF-NCR, is consistent with previous measles virus sequencing studies (7) and did not affect sequence concordance.

There are some limitations. First, the PCR workflow requires the use of six separate primer pools, which increases hands-on time and cost compared with lower-pool assays and may be perceived as cumbersome. However, this design was necessary to robustly achieve complete genome coverage, particularly across challenging regions such as the MF-NCR. In our evaluation, the six-pool structure was critical for minimizing amplification bias and dropout in these regions. A second limitation of the present study is the unequal representation of measles virus genotypes, with a greater number of genotype B3 samples than genotype D8 samples included in the assay accuracy assessment. Nevertheless, the primer set originally described by Zubach et al. 2025 (10) was extensively validated using genotype D8 strains. Modifications introduced in the current assay were designed to expand genotype coverage and did not alter the original D8-optimized primer sequences, supporting the robustness of the assay across both genotypes.

Overall, the Illumina-based tiled amplicon method described here provides a sensitive, accurate, and reproducible approach for measles virus whole-genome sequencing. By leveraging existing Illumina infrastructure, this workflow can be readily implemented in public health laboratories and supports enhanced genomic surveillance, outbreak investigation, and measles elimination efforts.

## Acknowledgments

We gratefully acknowledge our provincial partners for referring specimens to the NML for measles surveillance and genotyping. We would also like to acknowledge the Computational and Operational Genomics Section at the NML for bioinformatic and pipeline support.

