## Supplementary figures and images for "Measles Whole Genome Sequencing by an Illumina Tiled Amplification Method"

### Supplemental Figure

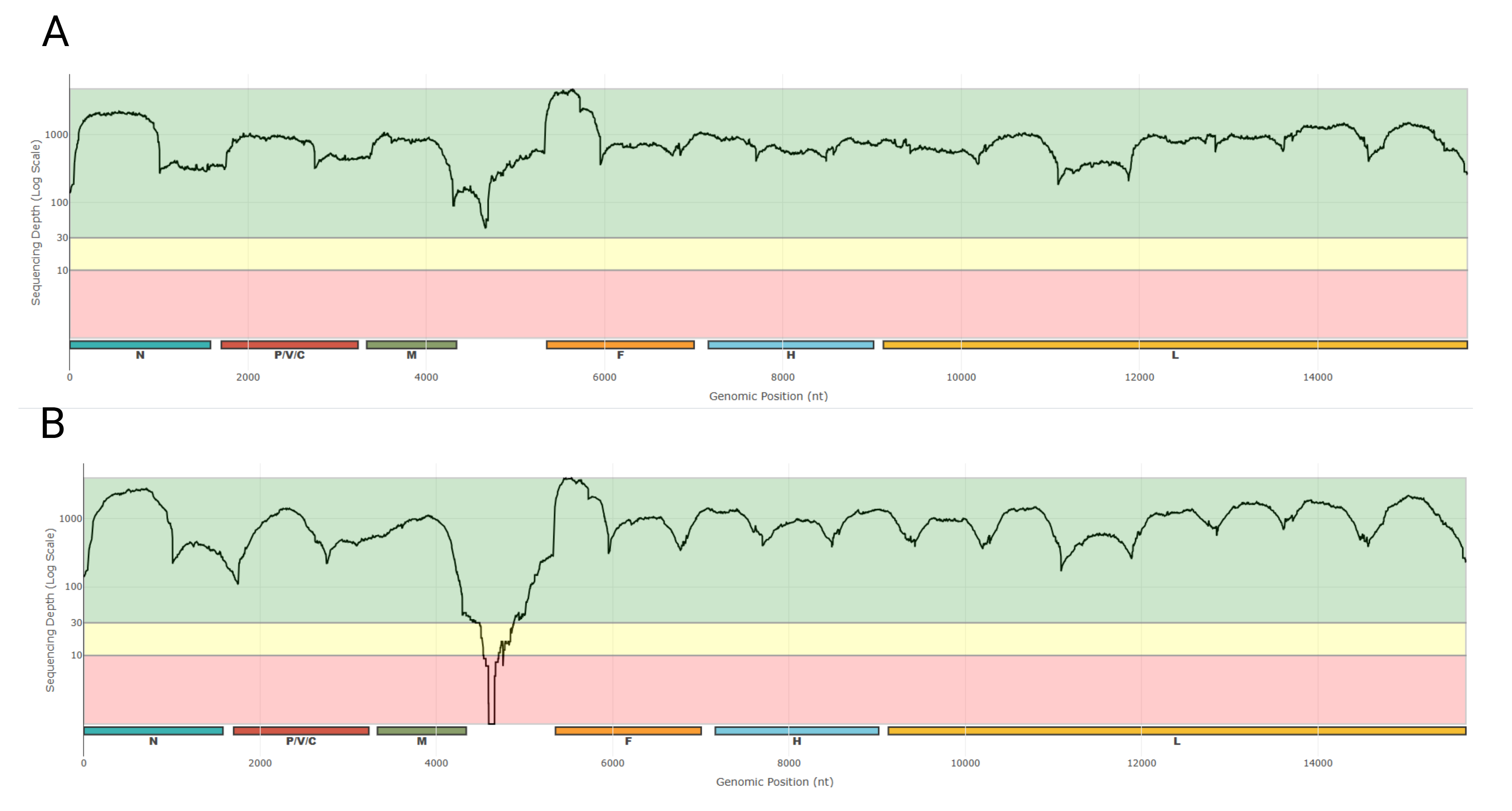
